# Shared Nucleotide Flanks Confer Transcriptional Competency to bZip Core Motifs

**DOI:** 10.1101/262659

**Authors:** Daniel M. Cohen, Hee-Woong Lim, Kyoung-Jae Won, David J. Steger

## Abstract

Sequence-specific DNA binding recruits transcription factors (TFs) to the genome to regulate gene expression. Here, we perform high resolution mapping of CEBP proteins to determine how sequence dictates genomic occupancy. We demonstrate a fundamental difference between the sequence repertoire utilized by CEBPs in vivo versus the palindromic sequence preference reported by classical in vitro models, by identifying a palindromic motif at less than 1% of the genomic binding sites. On the native genome, CEBPs bind a diversity of related 10 bp sequences resulting from the fusion of degenerate and canonical half-sites. Altered DNA specificity of CEBPs in cells occurs through heterodimerization with other bZip TFs, and approximately 40% of CEBP-binding sites in primary human cells harbor motifs characteristic of CEBP heterodimers. In addition, we uncover an important role for sequence bias at core-motif-flanking bases for CEBPs and demonstrate that flanking bases regulate motif function across mammalian bZip TFs. Favorable flanking bases confer efficient TF occupancy and transcriptional activity, and DNA shape may explain how the flanks alter TF binding. Importantly, motif optimization within the 10-mer is strongly correlated with cell-type-independent recruitment of CEBPβ, providing key insight into how sequence sub-optimization affects genomic occupancy of widely expressed CEBPs across cell types.

## INTRODUCTION

Sequence-specific DNA binding by transcription factors (TFs) is fundamental to the establishment and maintenance of gene programs that drive cell function in health and disease (1, 2). The genomic distribution of TFs at enhancers and promoters defines the framework by which these proteins orchestrate temporal and spatial regulation of gene expression (3, 4). The genomic landscape of TF-binding sites (TFBSs) is organized by the non-random distribution of DNA recognition sequences, or motifs, that mediate recruitment of their cognate TFs (5). Consequently, defining the motif preferences employed by each TF and mapping the genomic locations of motifs are key to unlocking the basis for gene regulatory networks.

High-throughput approaches have facilitated the identification of TFBSs both in vitro and in vivo (6–8). Protein binding microarrays (PBMs) and high-throughput in vitro selection (HT-SELEX) have determined the specificities of hundreds of isolated TFs from multiple species (9, 10). Alternatively, chromatin immunoprecipitation combined with next generation sequencing (ChIP-seq) has been employed extensively to locate where TFs occupy the native genome and to interrogate motifs from overrepresented sequences in ChIP-seq peaks. In spite of the high information content of consensus sequences, experimentally determined TF motifs have limited ability to predict in vivo binding (11). DNA accessibility (12–14) as well as contextualizing factors including DNA shape (15ߝ19), DNA methylation (20–22), neighboring TF interaction (23, 24) and altered sequence specificity due to heterodimer formation between related TFs (25) constrain and reshape how TFBSs are utilized in native genomes. Collectively, these variables help explain why TFs occupy only a small fraction of candidate motifs in the genome (26).

Contrary to the strong sequence dependence of TF binding in vitro, it has been suggested that TFs are recruited independently of their cognate sequences at many ChIP-seq peaks either through indirect protein-protein interaction (tethering) (24, 27–30) or through recognition of DNA shape (31, 32). Coupled with the observation that motif scores fail to differentiate between bound versus unbound genomic sequences (18, 33, 34), the question of what constitutes the minimal sequence determinants for TFBSs in vivo has become increasingly uncertain. Fortunately, new experimental approaches are providing avenues to address this question. High resolution (20-50 bp) mapping of bound genomic sequences has been facilitated by the development of ChIP with lambda exonuclease digestion and sequencing (ChIP-exo) (35). Close discrimination of bound motifs can revise and improve recognition sequences (36, 37) and resolve dimeric versus monomeric binding (38, 39). In parallel, comparison of bound and unbound motifs in biochemical assays of TF binding to histone-free genomic DNA is providing further insight into the native sites that are sufficient to mediate occupancy (40–43). Uniting these approaches has the potential to bridge major gaps in our understanding of the relationship between TF sequence specificity, motif occurrence and occupancy at native genomic sites.

CEBP TFs are particularly interesting in terms of how DNA-binding specificity defines genomic occupancy for two key reasons. As lineage determining TFs in several tissues (44–47), CEBPs may function as pioneer factors that overcome the inhibitory effects of chromatin, and thus defining their sequence specificity may be instructive as to whether a relationship exists between binding site affinity and TF occupancy in the genome. In addition, CEBPs can bind DNA as both homodimers and heterodimers (47), and their ability to target different sequence motifs through heterodimerization with other bZip family members (25, 41, 48–50) may enable the utilization of a broad repertoire of motifs to control a variety of gene expression programs (51). Indeed, CEBPs occupy tens of thousands of sites in primary cells and tissues (26, 52–54), however degenerate ChIP-seq motifs obscure the importance of sequence determinants for binding site selection.

Here, we report the high-resolution mapping of CEBP-binding sites in the human and mouse genomes using ChIP-exo. We find that CEBPs occupy a large repertoire of sequences in vivo defined by the fusion of canonical and degenerate CEBP half sites. Positive selection for the nucleotide composition observed within the core motif reflects altered sequence specificity of CEBP homo-versus heterodimers. We demonstrate the importance of the CEBP 10-mer motif by identifying an optimal sequence that is prevalently bound independent of cell type, suggesting that it forms a high-affinity-binding site that overrides chromatin context. Moreover, we reveal a critical role for negative sequence selection, i.e. the exclusion of a particular base, at the first and last position of the 10-mer. Negative selection for specific flanking nucleotide pairs is a general feature shared by multiple bZIP TF motifs, and the distinction between favorable and unfavorable motif contexts correlates with changes in DNA shape. We illustrate the functional importance of flanking base composition by showing that natural genetic variation from single nucleotide polymorphisms (SNPs) that introduce non-permissive flanking bases leads to strain-specific CEBP occupancy in mice. Collectively, these findings provide an expanded motif definition for CEBP that can be generalized to the broader bZip family, and establish important relationships between motif optimization, genomic occupancy and transcriptional activity.

## MATERIALS AND METHODS

Animal experiments were reviewed and approved by the Institutional Animal Care and Use Committees of the University of Pennsylvania. Mice were kept under standardized conditions with water and food *ad libitum* in a pathogen-free animal facility. hMSCs were obtained from Lonza and maintained in low glucose Dulbecco’s modified Eagle’s medium supplemented with 10% fetal bovine serum and 2mM glutamine. hMSCs were used at passages 4 through 7. ChIP-exo was performed as described previously (38) with minor modifications explained in the Supplementary Information. Extended and detailed methods for computational analyses of the genomics data are available in the Supplementary Information. Luciferase reporter assay and DNA pyrosequencing are described in the Supplementary Information.

## RESULTS

### CEBP proteins recognize a diversity of genomic sequences through a degenerate half site

To identify the genomic sequences targeted by CEBP TFs with high resolution in the native genome, we performed ChIP-exo in primary human mesenchymal stem cells (hMSCs) for CEBPβ as well as in mouse liver tissue for CEBPα and CEBPβ. The approach uses lambda exonuclease to trim ChIP DNA until a bound protein blocks further enzymatic activity (55). This creates 5’ borders on both DNA strands that are juxtaposed with the protein, manifested as opposite-stranded peak pairs on a genome browser, and achieves 20-50 bp resolution of DNA binding (35).

Opposite-stranded peak pairs annotate both canonical CEBPβ homodimer motifs and CEBPβ-sequences bound by the ATF4 heterodimer in hMSCs, demonstrating the resolving power of ChIP-exo (Figure 1A). Globally, CEBP peak pairs show an average distance distribution of 15-30 bp, with a predominant distance of 25 bp for CEBPβ and 27 bp for CEBPα (Figure 1B, Supplementary Figure S1A). Motif analysis reveals exclusive enrichment of an 8-mer-core sequence comprised of a degenerate half site (TKnn) fused to a CEBP half site (GCAA) (Figure 1C, Supplementary Figure S1B). Ordered peak pairs flank this motif at a majority of ChIP-seq peaks (Figure 1D, Supplementary Figure S1C), indicating that CEBPs occupy the genome primarily through direct, sequence-specific interaction. Parsing the CEBP cistrome by individual 8-mer variants of the CEBP core motif reveals that the sequence bound most frequently by CEBPβ and CEBPα is TTGTGCAA (Figure 1E), partly due to its high occurrence in the genome (Supplementary Figure S1D). Nevertheless, this sequence accounts for only about 14% of high-confidence ChIP-exo-annotated binding sites. Together with similar ChIP-seq occupancy strengths observed as a function of CEBP 8-mers (Supplementary Figure S1E), these data indicate that no singular sequence explains the majority of CEBP binding. Interestingly, the CEBPβ-ATF4 heterodimer sequence, TGATGCAA, is the second most prevalent CEBP core motif variant, and additional hybrid motifs composed of non-CEBP bZip half sites (TGWN) joined to the CEBP half site are also present within the top-ranked sequences. The tolerance of substituting G in lieu of the canonical T at the 2nd position of the hybrid core suggests either an intrinsic relaxation of CEBP’s sequence specificity in physiological contexts, broadened motif recognition through heterodimerization, or both. As a whole, the ChIP-exo data demonstrate conservation between human and mouse CEBP family members through interaction with a compound motif anchored by a CEBP half site.

**Figure 1.**
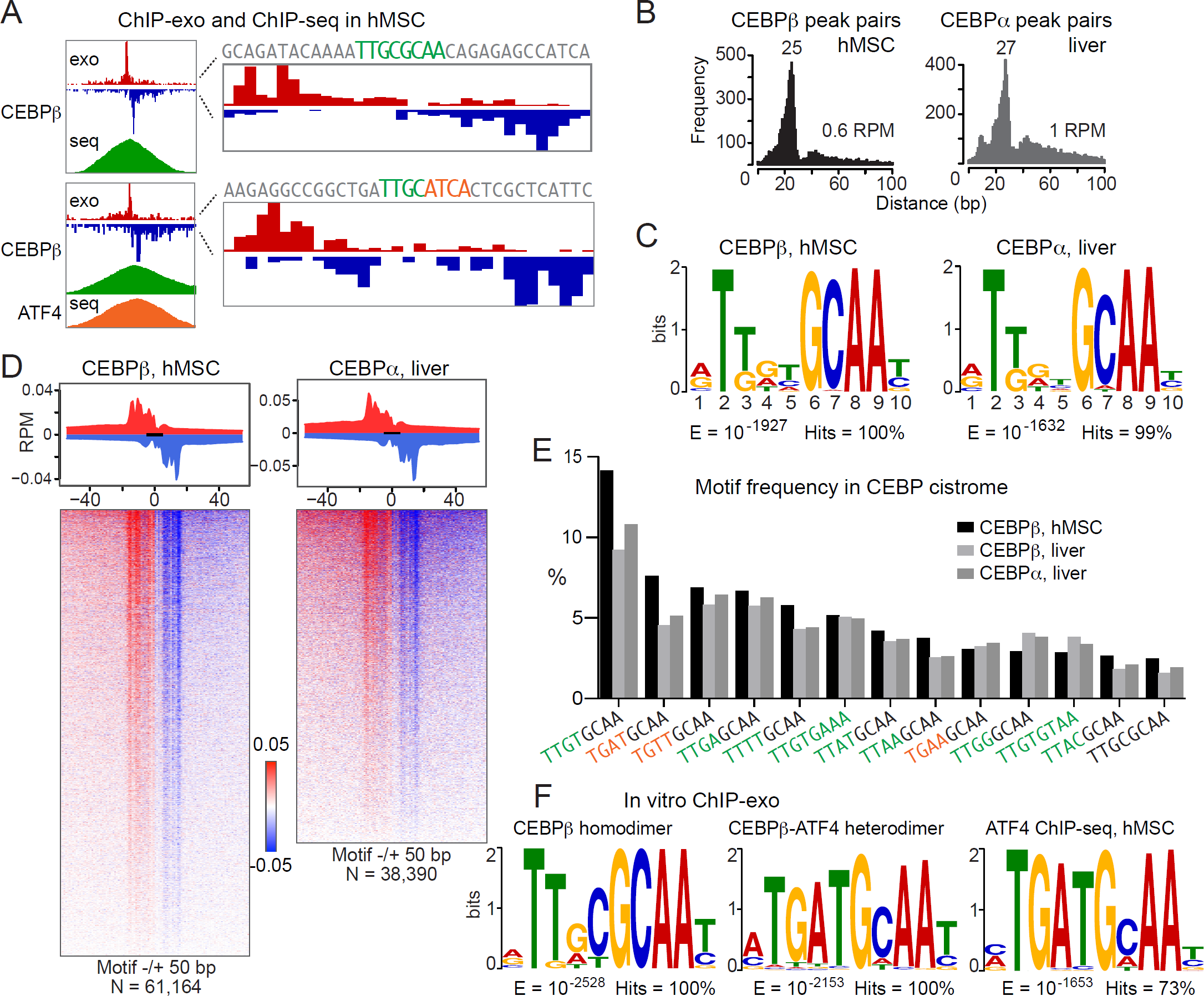
CEBP proteins occupy multiple sequence motifs on the native genome. (**A**) Comparison of ChIP-exo and ChIP-seq results for CEBPβ in hMSCs. Left, an opposite-stranded peak pair from ChIP-exo resides near the center of the ChIP-seq peak for either a homodimer-binding site (top) or a heterodimer site with ATF4 (bottom). Right, closer inspection reveals canonical DNA motifs for CEBPβ (green) or CEBPβ-ATF4 (green-orange) between the ChIP-exo peak pairs. Red and blue indicate the 5’ ends of the forward- and reverse-stranded sequence tags, respectively. (**B**) Distance distributions for the spacing between opposite-stranded peak pairs. Predominant distances are indicated. (**C**) MEME de novo motif analyses of the 1000-top-ranked ChIP-exo peak pairs spaced 15-30 bp apart. (**D**) Average profiles (top) and density heat maps (bottom) of the ChIP-exo sequence tags at CEBP-binding sites in hMSCs or liver. (**E**) Top-ranked core motifs at CEBPβ peak pairs in hMSCs compared with CEBPs in liver. The CEBP half site, GCAA, is uncolored; degenerate half site is green (CEBP related) or orange (bZip related). (**F**) MEME de novo motif analyses of the 1000-top-ranked peak pairs spaced 10-30 bp apart are shown for the CEBPβ homodimer and CEBPβ-ATF4 heterodimer. Motif analysis from ATF4 ChIP-seq in hMSCs is shown for comparison.

The CEBP motif identified in primary cells and tissue differs strikingly from the optimal sequence observed for homodimers in vitro. Both early studies (56–59) and more recent systematic biochemical approaches (9, 10, 60) report that the CEBP homodimer binds a palindromic motif formed by the fusion of two CEBP half sites (ATTGCGCAAT), yet this sequence only represents only a small fraction (≤ 1%) of the genomic sites occupied by CEBPs in vivo. To exclude the possibility that this discrepancy is caused by assay-dependent effects, we performed ChIP-exo utilizing recombinant CEBPβ homodimer or ATF4-CEBPβ heterodimer and protein-free genomic DNA. A sequence resembling the palindromic CEBP motif is enriched at peak pairs for the CEBPβ homodimer (Figure 1F), consistent with findings from PBMs (10), HT-SELEX (9) and SMiLE-seq (60). Yet, this motif is distinct from the consensus motif for endogenous CEBP. In contrast, in vitro ChIP-exo for the CEBPβ-ATF4 heterodimer yields a motif that is very similar to that reported for ATF4 in hMSCs (Figure 1F) (41). Sequence-specific interaction by the CEBPβ homo- and heterodimer is indicated by the emergence of peak pairs with fixed spacing that flank both motifs (Supplementary Figures S1F and S1G). Thus, in vitro ChIP-exo corroborates the DNA sequence specificity reported by established biochemical approaches, and illustrates a fundamental difference between the DNA-binding specificity of the CEBPβ homodimer versus CEBPβ in cells. Although a few thousand heterodimeric sites of ATF4 with various CEBPs have been mapped in vivo (41, 48, 50), they represent only 2-5% of the CEBPβ cistrome in hMSCs. Thus, the observation that roughly 40% of CEBPβ binding in hMSCs occurs at hybrid sequences comprised of AP-1 or ATF-like half sites fused to a CEBP half site suggests that heterodimerization with other bZip family members may occur on a much broader scale than previously envisioned.

### Sequence optimization regulates cell-type-specific binding by CEBPβ

Despite the strong preference of CEBP homodimers for the palindromic sequence, TTGCGCAA, it accounts for less than 3% of all CEBPβ binding sites. However, correcting for the fact that the CEBP palindromic 8-mer occurs rarely in the human genome, it exhibits the highest fraction of genomic occupancy of any CEBP motif (Supplementary Figure S1D). This suggests a relationship between sequence optimization and probability of genomic occupancy. To test whether motif optimization is correlated with the likelihood of binding and/or transcriptional regulation, we examined the relationships between CEBPβ-binding sites, CEBP motifs, and gene transcription across multiple human cell lines (hMSCs, Helas3, HepG2, K562, IMR90, A549). Consistent with frequency measurements for shared versus unique binding sites for a TF in different cell types (27, 61), and CEBPβ specifically (62), approximately 20% of CEBP sites map to either a single cell type (cell-type specific) or all cell types (cell-type independent), respectively, while the remaining 60% fall between these extremes (Figure 2A). Functional CEBPβ sites show enrichment for RNA Polymerase II (RNAPII) (62), and using gene body RNAPII occupancy as a surrogate for transcription, we observed higher transcriptional activity at genes near cell-type-independent sites compared to genes near cell-type-specific sites (Figure 2B). Moreover, distinct gene ontologies were observed, with genes near cell-type-independent sites enriched for general processes such as mRNA metabolism and translation, whereas cell-type-specific site-gene pairs associate with specialized pathways such as adipocytokine signaling and lipoprotein metabolism. Intriguingly, the genomic distribution of cell-type-independent sites is biased towards transcription start sites (TSSs), whereas cell-type-specific sites display a gene-distal distribution characteristic of the overall CEBPβ cistrome (Supplementary Figure S2A).

**Figure 2.**
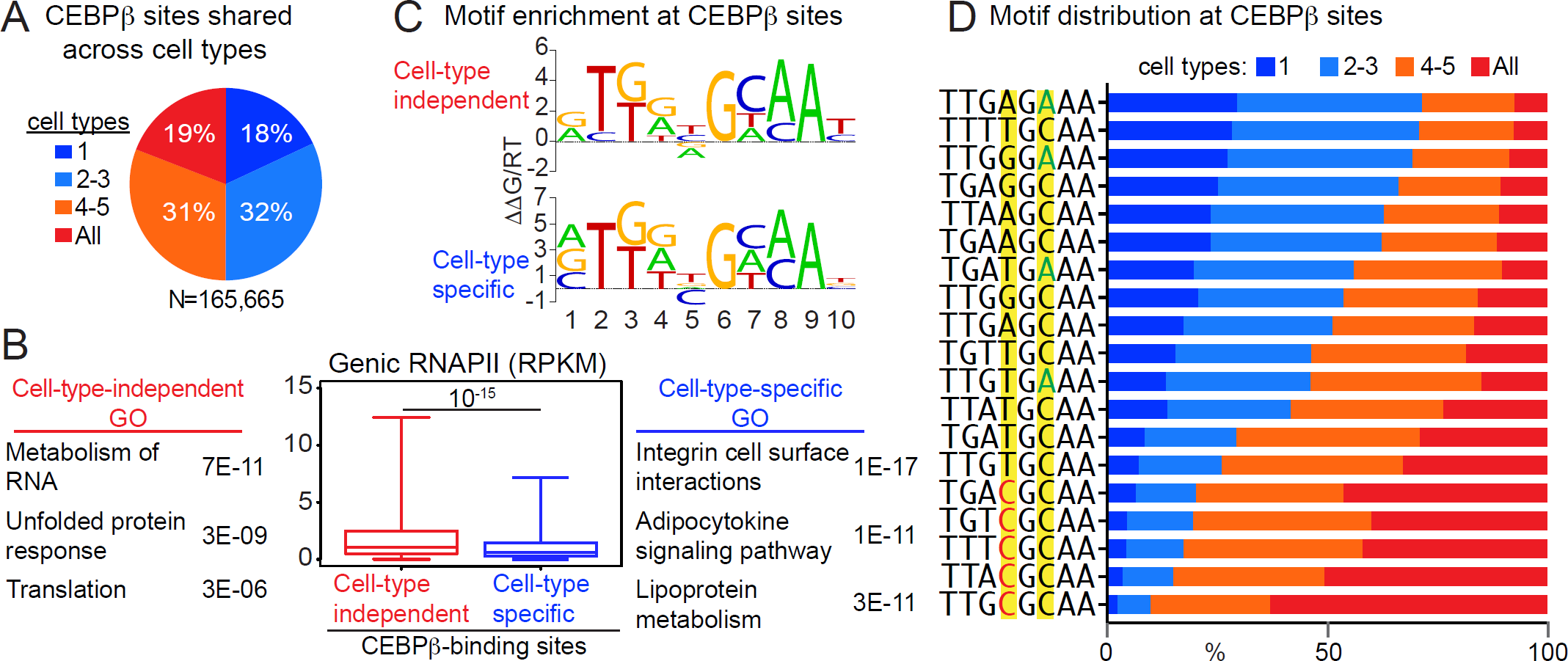
Selective versus widespread occupancy across cell types for distinct CEBPβ motifs. (**A**).Pie chart comparing CEBPβ occupancy across 6 human cell types (hMSC + 6 ENCODE cell lines). sites unique to one cell type (cell-type-specific) or shared between 2-3, 4-5, and all 6 cell types. (**B**) Box plot interrogating RNAPII occupancy (ChIP-seq reads per million, RPM) at expressed genes within 100 kb of cell-type-independent or cell-type-specific binding sites for CEBPβ. Wilcoxon rank sum test used to compare adjacent classes. Highly ranked gene ontology (GO) terms are shown. (**C**) De novo motif analyses showing de-enriched bases that differ between cell-type-independent and cell-type-specific sites. (**D**) Individual 8-mers enriched at cell-type-independent or cell-type-specific sites for CEBPβ were examined to display occupancy across the 6 human cell types. Base positions of interest within the first and second half sites are highlighted.

Motif quality has been reported to correlate with shared occupancy across cell types for some nuclear receptors (27, 63), but prior studies of cell-type-dependent binding for CEBPβ focused on the role of collaborating TFs and chromatin (62). Interestingly, de novo motif analyses revealed a depletion of C in the 5^th^ position of the CEBP motif at cell-type-specific, but not cell-type-independent, sites (Figure 2C). This C conforms to the canonical CEBP half site, suggesting that the probability of CEBPβ occupancy is correlated with motif optimization. To further elucidate this relationship, we surveyed the utilization of the top motif variants in both classes of sites. Preservation of C at the 4^th^ position of the core 8-mer (5^th^ position of the 10-mer) was strongly correlated with increased probability of binding in multiple cell types, with 90% co-occupancy of the CEBP palindrome in 4 or more cell types (Figure 2D). Conversely, cell-type-specific sites are enriched for sub-optimized motifs, harboring substitutions in the 4^th^ and 6^th^ positions of the core 8-mer. Consistent with the hypothesis that cell-type-independent binding sites are privileged for both high-affinity motifs and highly accessible chromatin, average CEBPβ-binding strength is positively correlated with co-occupancy status (Supplementary Figure S2B). These data demonstrate that motif optimization within the core 8-mer increases the probability that any given CEBP-binding site will be shared across cell types, and also suggest a relationship between optimized CEBP motifs and increased transcriptional activity. Though rare genome-wide, optimized CEBP motifs may serve as elite sequences that enable CEBPβ to overcome the repressive effects of chromatin to coordinate a limited program of constitutive gene expression.

### Bases directly abutting the core CEBP motif impact occupancy

We would predict occupancy at that nearly every genomic instance of the palindromic CEBP core 8-mer if this motif constitutively recruits CEBPs, and yet most are unoccupied in hMSCs (Supplementary Figure S1D). To investigate potential distinguishing features for bound versus unbound CEBP palindromes, we profiled CEBPβ occupancy at all 3121 palindromic 8-mers (excluding unplaced contigs) in the human genome using our in vitro ChIP-exo data derived from recombinant CEBPβ homodimer bound to histone-free genomic DNA. While the vast majority (84%) of CEBP palindromes showed binding, a subset failed to recruit CEBPβ. Sequence alignments of these unbound sites revealed a pronounced difference in the base composition at positions immediately flanking the core motif (Figure 3A). In contrast, neighboring positions beyond these flanks have random sequence variation (Supplementary Figure S3A). Strikingly, the occurrence of T at the 5’ flank or A at the 3’ flank is negatively correlated with CEBPβ occupancy. Moreover, while C is also disfavored at the 5’ flank, its ability to cripple the functionality of the palindromic 8-mer is most pronounced when paired with A at the 3’ flank. Likewise, T-G dinucleotide flanks appear highly deleterious to CEBPβ binding (note that **C**TTGCGCAA**A** and **T**TTGCGCAA**G** are reverse-complementary 10-mer sequences). Importantly, sequence preferences at these flanking positions are implicit within the motif logos from our ChIP-exo experiments (Figure 1C, Supplementary Figure S1B). Careful inspection of these logos reveals the exclusion of T at the 5’ flank and A at the 3’ flank. However, compared to the core 8-mer, the relatively low information content encoded in these flanks de-emphasizes their importance, and fails to underscore how specific base pairings at the 5’ and 3’ flanks (T-A, C-A, T-G) can override the ability of an optimized 8-mer core sequence to recruit CEBPβ.

**Figure 3.**
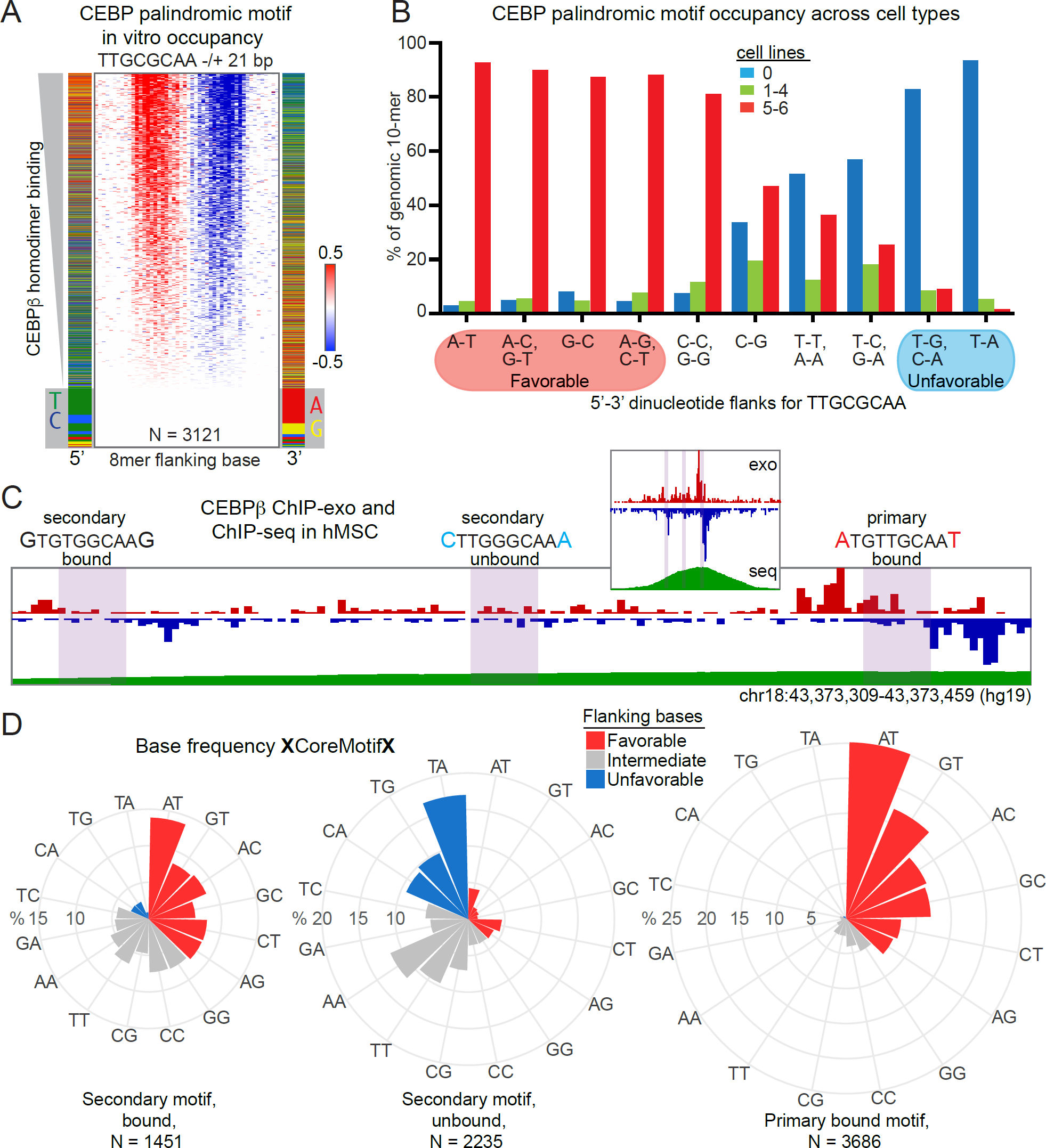
Bases directly flanking the core CEBP 8-mer affect occupancy. (**A**) Density heat map of the in vitro ChIP-exo reads for the CEBPβ homodimer at all canonical palindromic 8-mers with mappable sequence. Binding strength is ordered from top to bottom. Color charts show the base identity at the first position next to the 8-mer on the 5’ and 3’ ends. Grey boxes indicate sites without detectable ChIP-exo reads. (**B**) Histogram interrogating relationship between flanking bases and CEBPβ occupancy at all canonical palindromic 8-mers across 7 cell types. Equivalent flanking pairs (5’-3’) are grouped together. ENCODE ChIP-seq peak calls are plotted. Favorable flanks associate with occupancy in most cell types at most locations. Unfavorable flanks are not bound in any cell type at most locations. (**C**) CEBPβ ChIP-exo reads at a ChIP-seq peak (insert) from hMSCs with 3 CEBP motifs of the form TKnnGCAA. Purple shading indicates motif locations. Binding to the primary motif (right) is indicated by co-localization with an opposite-stranded peak pair. The secondary motifs co-localize with either a weaker peak pair having too few reads to meet binding cutoffs (left) or background reads (center). Flanking bases (larger font) are indicated as favorable (red) or unfavorable (blue) based on the findings from the palindromic 8-mer. (**D**). Polar bar graphs indicating the frequency of each pair of bases (5’ and 3’) flanking a generic CEBP motif at primary and secondary motifs. Comparison of the frequencies for favorable, intermediate and unfavorable flanks between the secondary weakly bound versus unbound motifs are highly statistically significant (*p* < E-13 by hypergeometic distribution).

The apparent importance of the flanking bases indicates that the minimal sequence determinant for CEBP binding to the genome is encoded by a 10-mer sequence, corroborating earlier biochemical and structural studies that identified a 5-mer half-site (56–59). To further investigate the relationship between sequence variation at core-motif flanking positions and motif binding strength, we revisited our analysis of cell-type-dependent occupancy across all 3121 CEBP palindromic motifs, parsing by 10-mer variants (Figure 3B). Remarkably, 88% of all genomic instances of the top-6 performing 10-mers are co-occupied by CEBPβ in 5 or more cell types (red bars), and > 95% of these sequences are bound in at least one cell type (sum of red and green bars). Conversely, 10-mers comprised of a palindromic 8-mer nested within unfavorable flanks are rarely occupied. These data indicate that flanking nucleotides play a critical role in CEBP motif recognition, and reveal that when annotated as a 10-mer sequence, the CEBP palindrome functions to recruit CEBPβ independently of cell type or genomic neighborhood. In contrast, 8-mer palindromic sequences with neutral flanking dinucleotide pairs (combinations of one favorable and one unfavorable flanking nucleotide at the 5’ and 3’ positions) are enriched for cell-type-specific CEBPβ binding, suggesting that while these sequences have the potential to recruit CEBPs, they are more sensitive to cell-type-specific differences in chromatin structure. Reduced affinity for these sequences could explain this behavior, which is indicated by weaker ChIP-exo signal at sites with neutral flanking bases relative to those with favorable flanks (Supplementary Figure S3B).

These findings enhance our understanding of the optimal CEBP palindromic motif, yet an important question is whether core-motif flanking nucleotides are also important in the context of a more degenerate CEBP motif that is representative of the broader CEBP cistrome. However, the very nature of these degenerate sequences as sub-optimized binding sites presents a challenge in interpreting whether unbound genomic sequences fail to recruit CEBPβ due to chromatin effects, unfavorable flanks, or both. Importantly, the effects of chromatin can be excluded by testing how CEBP 8-mers are populated within known CEBPβ ChIP-seq peaks, which by definition reside within accessible chromatin. While ChIP-exo has the resolution to resolve multiple binding events within single ChIP-seq peaks for CEBPβ, our analysis pipeline picks the strongest peak pair with a characteristic spacing per ChIP-seq peak, and thus does not explicitly identify every instance of a bound motif. As a result, we re-interrogated our ChIP-exo data to discover additional, weaker CEBPβ-binding events within CEBPβ ChIP-seq peaks, and compared their motifs to co-localized 8-mer sequences that failed to exhibit a characteristic ChIP-exo peak pair. A total of 3686 CEBP motif candidates were mapped in the vicinity of a CEBP-bound motif, and classified into bound versus unbound sequences based on their ChIP-exo signature (Figure 3C, Supplementary Figure S3C). Within the set of candidate secondary motifs, 61% lacked appropriately spaced opposite-stranded peak pairs, indicating little or no occupancy. We then examined the frequency of dinucleotide flanks abutting the bound primary and secondary CEBP 8-mers as well as the unbound secondary 8-mers (Figure 3D). Consistent with the behavior of the flanks surrounding the palindromic 8-mer, the flanks present at primary (strong) ChIP-exo bound motifs are enriched for A-T, A-C, A-G, G-T, G-C, and C-T dinucleotide flanks, whereas the unfavorable T-A, C-A, and T-G flanks are essentially absent. This frequency distribution differs from the background rate for CEBP 8-mers across the human genome (Supplementary Figure S3D), such that favorable flanks occur less frequently than expected by chance. Secondary bound CEBP motifs show a preference for favorable flanks that is similar to primary sites, albeit at lower frequencies. In contrast, unbound 8-mers are enriched for unfavorable and neutral flanks. These data demonstrate that flanking bases play a role in discriminating which candidate CEBP 8-mers are bound within regions of open chromatin. Combined with the earlier analyses, they reveal that a 10-mer sequence enables discrimination of real versus decoy CEBP motifs in the genome.

### Bases directly abutting core bZip motifs affect DNA shape

Crystallography studies of CEBPα and several other bZip proteins complexed with DNA have modeled contacts to the core bases (64–68), whereas interaction with the DNA backbone may take place at the flanks (67). These observations challenge the notion that the flanks interact with CEBPs via base readout, and led us to consider whether the flanking bases could impact genomic occupancy through alteration of DNA shape. To address DNA shape in an unbiased manner, we examined the CEBP motifs residing in accessible chromatin of hMSCs yet differing in their ability to recruit CEBPβ (see Supplementary Figure S3C). Comparison of intrinsic DNA shape features revealed significant differences between bound versus unbound motifs (Figure 4A). The shape changes occurred at or near the flanking bases, consistent with an intimate association between sequence and shape, and suggesting that CEBPβ prefers to bind motifs surrounded by more positive roll, less helical twist, wider minor groove width, and less negative propeller twist.

**Figure 4.**
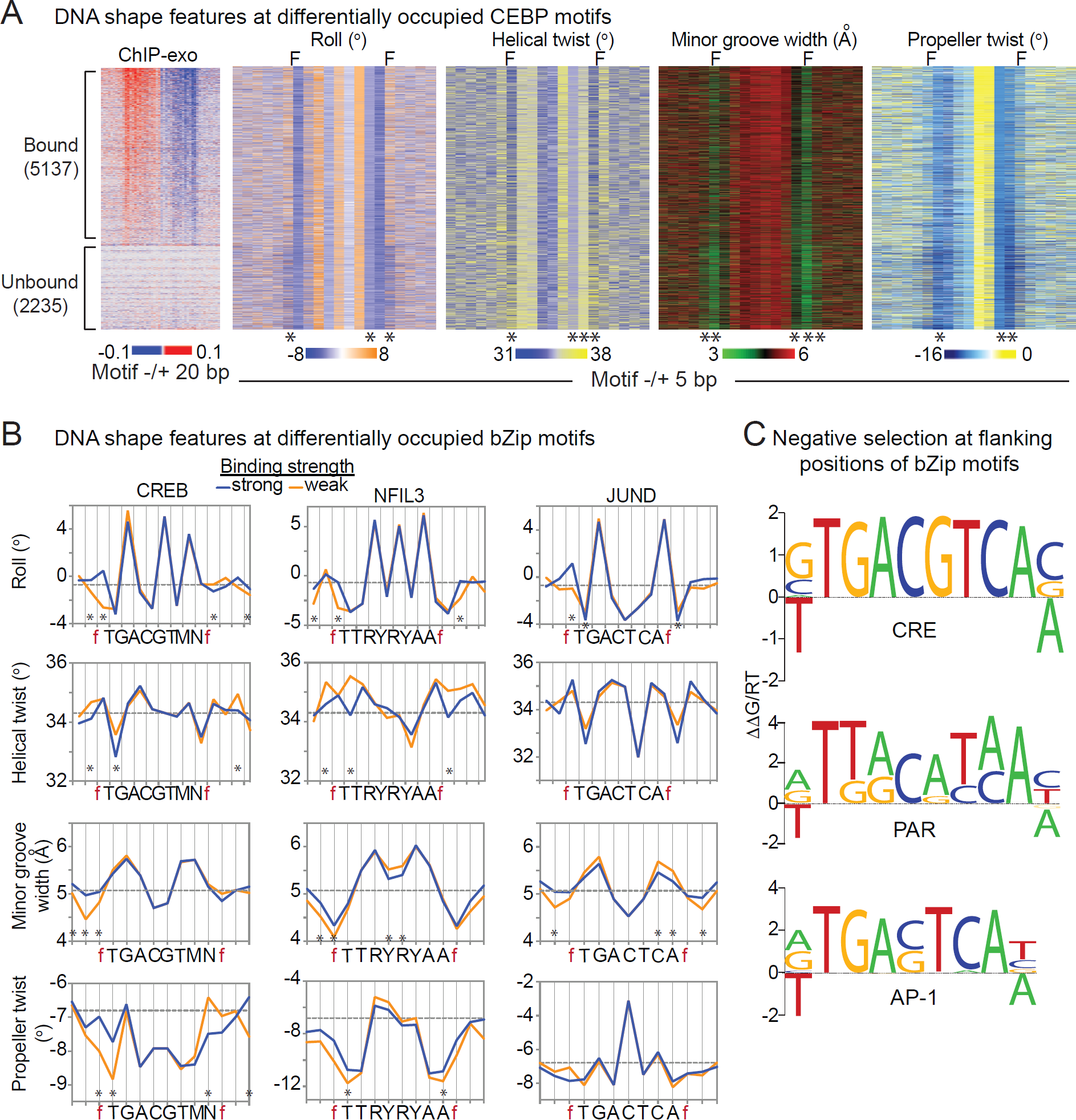
Bases directly flanking core bZip motifs influence DNA shape. (**A**) Density heat maps of DNA shape features (roll, helical twist, minor groove width and propeller twist) at occupied and unoccupied CEBP motifs within CEBPβ ChIP-seq peaks from hMSCs. CEBPβ ChIP-exo signal sets the ordering for all heat maps. F indicates the positions flanking the core 8 bp motif. * denotes p < E-50, Wilcoxon rank sum test. (**B**) Average profile plots of DNA shape parameters at weakly and strongly bound motifs for CREB, NFIL3 and JUND. Differential binding was determined by protein binding microarrays (10). Core motifs and flanking positions (f) are shown. Dashed line indicates the genome-wide average for each shape feature. *, denotes p < 0.01, Wilcoxon rank sum test. (**C**) Motif analyses of CREB1, NFIL3 and AP-1 ChIP-seq data. Top-ranked motif is shown for each emphasizing de-enriched bases in the 5’ and 3’ positions flanking the core sequences. Negative selection of bases within the cores is not shown.

More broadly, we sought to determine whether the DNA shape features associated with CEBPβ occupancy apply to bZip TFs in general. To examine a potential relationship between DNA shape and bZip TF occupancy, we identified weakly and strongly bound motifs from published PBMs for CREB, NFIL3 and JUND (10). Comparison of differentially bound motifs for each TF revealed significant alterations for various shape features (Figure 4B). Interestingly, the shape changes resembled those at the CEBP motif, suggesting that CREB, NFIL3 and JUND prefer to bind motifs surrounded by more positive roll, less helical twist, wider minor groove width, and less negative propeller twist. Given that DNA sequence determines shape, we examined the motifs enriched in previously published ChIP-seq datasets for CREB, NFIL3 and AP-1 (69). Similar to the CEBP motif, we found a clear exclusion of T and A in the 5’ and 3’ flanking positions, respectively (Figure 4C). Thus, the motifs for multiple bZip proteins share similar negative selection for T-A pairs at the flanks, indicating that an expanded motif definition including flanking bases is broadly important across mammalian bZips.

### Bases directly abutting core bZip motifs affect transcriptional activity

To test whether a change in the flanks is sufficient to convert a functional CEBP-binding site into a crippled site, we performed ChIP for CEBPs in liver tissue isolated from C57BL/6J (B6) and 129S1/SvImJ (129) mice, and examined occupancy at sites carrying SNPs that introduce unfavorable flanks into CEBP-binding sites when comparing B6 to 129. While only two sites exist meeting the criteria for type of nucleotide substitution of interest and the absence of a neighboring CEBP-binding site, both show diminished occupancy of CEBPα and CEBPβ in 129 mice relative to B6 (Figure 5A). Consistent with these results, B6×129 F1 mice showed significantly skewed binding of CEBPs to the B6 alleles (Figure 5B). Because the B6 and 129 alleles reside in the same nuclei of F1 mice and are thus exposed to the same trans-acting factors, these data demonstrate that cis effects determine differential binding of CEBPs at these loci. Specifically, the introduction of unfavorable flanking nucleotides may be sufficient to impair CEBP binding independently of the core 8-mer.

**Figure 5.**
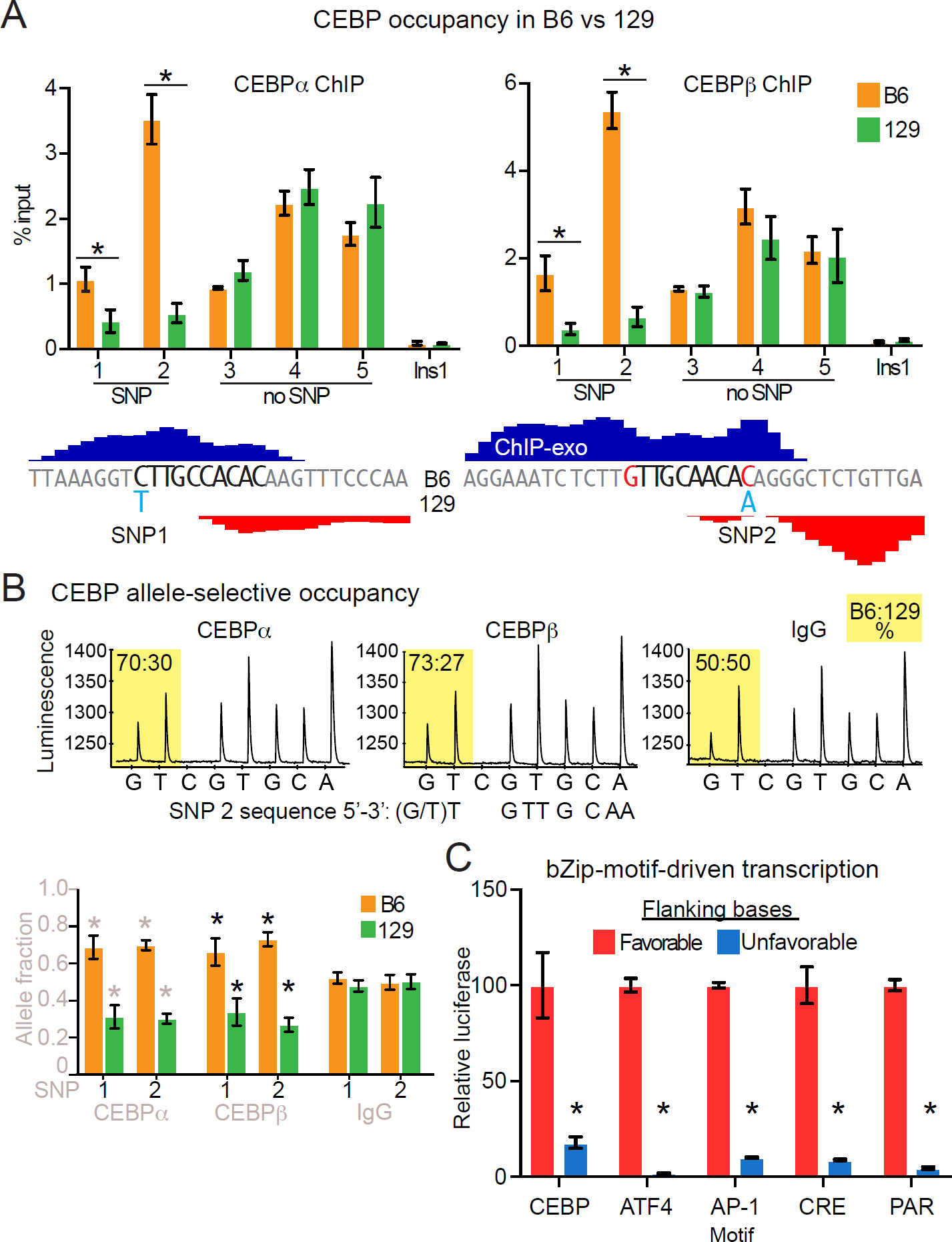
Bases directly flanking core bZip motifs regulate function. (**A**) CEBP ChIP in liver tissue isolated from B6 and 129 mice interrogating binding sites with and without SNPs in the bases flanking the core CEBP 8-mer. ChIP-exo tracks (bottom) show location of SNP relative to core 8-mer and opposite-stranded peak pairs. Ins1, non-specific control site. Error bars depict SEM from 5 biological replicates. *, denotes p < 0.05, Student’s *t*-test comparison of B6 with 129. (**B**) Pyrosequencing of CEBPα, CEBPβ and IgG ChIP DNA prepared from liver tissue of B6×129 F1 mice. Chromatograms show raw data for SNP2. Note that these data report the opposite DNA strand shown in A. Bar plot (lower left) reports results for SNPs 1 and 2 with error bars depicting SEM from 5 biological replicates. *, denotes p < 0.05, Student’s *t*-test comparison of CEBPα or CEBPβ with IgG. (**C**).Core bZip motifs were assembled into repeats of four and assayed by a luciferase reporter in HEK293T cells. Flanking bases (X-X) for the CEBP (XTTGTGCAAX), ATF4 (XTGATGCAAX), AP-1 (XTGACTCAX), CRE (XTGACGTCAX) and PAR (XTTACGTAAX) motifs were either favorable (A-T) or unfavorable (T-A, excepting the ATF4 motif, where a T-T pair was used to selectively target the flank of the ATF4 half site) for TF occupancy. Error bars depict SEM from 3 replicates. *, denotes p < 0.01, Student’s *t*-test comparison of favorable with unfavorable flanks for each motif.

Our data establish genome-wide trends between bZip TF occupancy and core-motif flanking bases. To test this relationship and extend its relevance to transcriptional activity, we examined the activity of synthetic luciferase reporters containing multimerized core motifs for distinct bZip TFs flanked by either favorable or unfavorable bases. Replacement of favorable with unfavorable flanks decreased luciferase activity across all bZip reporter constructs tested, with reductions ranging from 6-fold for the CEBP motif to ≥ 10-fold for the remaining motifs (Figure 5C). Thus, the data reveal a shared requirement across the bZIP family for favorable motif flanks that confer binding and transcriptional competency to their cognate core recognition sequences.

## DISCUSSION

We have used ChIP-exo to perform a genome-wide cataloging of motif utilization within the CEBP cistromes of primary human cells and mouse liver tissue. We demonstrate species-conserved sequence requirements for the recruitment of CEBP proteins to the native genome that are fundamentally different from the sequence preferences of CEBP homodimers in vitro. Pioneering in vitro studies (56–59) and more recent systematic biochemical approaches (9, 10, 60) report that the CEBP homodimer prefers to bind a palindromic motif formed by the fusion of two CEBP half sites (ATTGCGCAAT). Our data reveal that this motif captures less than 1% of the binding sites occurring in cells. On the native genome, CEBPs bind a diversity of related sequences resulting from the fusion of degenerate and canonical CEBP half sites that yields a 10-mer-consensus of the form VTKNNGCAAB. A large majority of binding sites, 70-90% depending on threshold cutoffs, contains bound CEBP motifs. This suggests that CEBPs primarily occupy the genome through direct, sequence-specific interaction, whereas binding to motifs with atypical spacing between half sites (25) and to other DNA-bound TFs through tethering contribute minimally to the genomic recruitment of CEBPs.

It is noteworthy that roughly 40% of CEBP-binding sites contain a G at the third position of the 10-mer, creating a preferred half-site motif for the ATF, AP-1, and CREB families of bZip TFs. Evidence has been found for the heterodimerization and altered sequence specificity of CEBP-ATF complexes compared to their homodimer counterparts (25, 41, 48–50), yet none of these studies addresses the extent to which heterodimerization drives genomic occupancy in vivo. For example, the identification of approximately 1600 ATF4-CEBPβ heterodimer sites in hMSCs represents only 2-5% of the total genomic CEBPβ sites (41). Our data indicate heterodimer binding greatly exceeding that with ATF4 such that CEBP occupancy of hybrid motifs represents a large fraction of the cistrome in vivo. This widespread occupancy is unprecedented and highly impactful for understanding the function of bZip TFs, and may help to explain why CEBPs populate large cistromes comprised of tens of thousands of binding sites in mammalian tissues and primary cells (26, 52–54).

Genome-wide cataloging of motifs within CEBP cistromes preserves genomic information that affords comparisons between bound and unbound sites. Direct examination of optimal TTGCGCAA 8-mers unoccupied both in vitro and in vivo identified T-A flanking bases that are disfavored for CEBPβ binding and transcriptional activation. Reminiscent of an early in vitro study of CEBPs (58), our finding impacts the understanding of DNA-binding specificity for bZip proteins in general, as the same flanking bases also cripple the transcriptional activity of the core motifs for ATF4, AP-1, CREB and PAR TFs. Negative selection against unfavorable flanks suggests that these positions contribute to motif recognition by modulating DNA-binding affinity. Structural studies of bZip proteins bound to DNA show interaction with the DNA backbone (67) but not the bases at the flanking positions (64–68), suggesting that the flanks impact bZip affinity through DNA shape affects. Contrasting the shape of bound and unbound sites for multiple TFs has led to the notion that motif flanks can regulate genomic occupancy through alteration of DNA shape (15, 18, 33, 70). Consistent with this, DNA shape features differ in similar ways between high- and low-affinity binding sites for CEBPβ, CREB, NFIL3 and JUND. The shared sequence bias at the flanks across distinct bZIP motifs, coupled with the fact that mono and di-nucleotide sequences account for more than 90% of the variance in commonly interrogated shape features (71), explains how similar DNA shape features can persist at the motif periphery even while divergent sequences dominate at the core.

Elite CEBP 10-mer motifs comprised of RTTRCGCAAY recruit CEBPβ in a cell-type-independent manner and are associated with higher levels of gene expression relative to cell-type-specific sites. Moreover, for optimized CEBP 10-mers containing a palindromic core, approximately 80% of genomic instances are bound by CEBPβ. Thus, highly optimized CEBP motifs are sufficient to recruit CEBPβ regardless of the genomic context, implying that CEBPs can overcome chromatin-mediated repression. Neutral flanks pair a favorable and unfavorable base at the first and last position of the 10mer, and they are correlated with a progressive loss of palindromic occupancy across cell types and weaker binding strengths in vitro. Importantly, these relationships between flanking sequence and motif occupancy can be generalized to the more degenerate CEBP motif, suggesting that CEBPs can populate lower-affinity sequences that are readily accessible in open chromatin.

Unlike high throughput assays that select for optimal TF-binding sequences, analysis of bound TF sequences suggests that optimized motifs play limited biological roles in genomic recruitment of TFs. A relationship between sub-optimized motifs and cell-type-dependent binding has been documented for ERα (27, 63), yet whether deviations from consensus motifs are biological drivers of differential TF occupancy is unknown, especially given the dominant effect of chromatin structure on the accessibility of DNA motifs. Intriguingly, motif sub-optimization through somatic mutation of the central CG dinucleotide of the CEBP motif has been reported in human cancers (72), suggesting an evolutionary pressure selecting against optimized CEBP motifs that mirrors the overall sparsity of these motifs in the human genome. Placed in the context of our work, perhaps the rarity of fully optimized TF motifs in eukaryotic genomes serves to limit constitutive genomic recruitment, suppressing the potential for TFs to trigger unregulated gene expression with regard to tissue or cell type. Conversely, the majority of CEBP-bound motifs are sub-optimized and occupied in a cell-type-specific manner. This observation fits with an emerging paradigm whereby tissue-specific gene expression is mediated by composite enhancers (24, 73–75) that recruit multiple TFs through sub-optimized motifs (73, 74). Rather than a fortuitous event, sub-optimization may be biologically favorable to impart a dependency of TF occupancy on chromatin environment, and render enhancers readily amenable to evolutionary turnover (76–78).

## ACCESSION NUMBERS

High throughput sequencing data have been deposited at GEO under accession number GSE111515.

## ACKNOWLEDGMENTS

We are grateful to Raymond Soccio for guidance on the experimental strategy interrogating strain-specific TF binding, and for providing liver tissue from 129S1/SvImJ and 129S1/SvImJxC57BL/6J F1 mice. We also thank Chris Krapp and Marisa Bartolomei for help with pyrosequencing and generously providing access to their sequencer. We are indebted to members of the Lazar laboratory for insightful discussions, and also thank the Functional Genomics Core of the Penn Diabetes Center (DK19525) for deep sequencing.

